# Induced-hyperandrogenism in female rats: assessing possible modulation of neurobehavioural and neurotransmitter changes following clomiphene/letrozole intervention

**DOI:** 10.1101/2023.07.16.549229

**Authors:** Olakunle J. Onaolapo, Olufemi O. Aworinde, Anthony T Olofinnade, Adejoke Y. Onaolapo

## Abstract

Hyperandrogenism is the excessive production of androgenic hormones resulting in infertility in a number of women. While letrozole and clomiphene citrate have been used to increase chances of achieving pregnancy, their effects on the brain has been scarcely studied. This study examined the effects of clomiphene and letrozole alone or in combination on neurobehavioural and neurochemical changes in female rats exposed to testosterone. Weaned rats were assigned into eight groups of ten each. Animals were grouped as normal control administered vehicle (normal saline) orally at 10 ml/kg or subcutaneously at 2 ml/kg, three groups administered clomiphene (CLOM) at 100 µg/kg, letrozole (LETR) at 5 mg/kg and or a combination of clomiphene and letrozole (CLOM/LETR) orally and saline subcutaneously. There were also four groups Testosterone (Test), Test/CLOM, Test/LETR or Test/CLOM+LETR administered testosterone enantate subcutaneously at 1 mg/100 g. Testosterone or saline was administered from day 1-35, while beginning on day 36, clomiphene, letrozole or saline was administered daily for 10 days. At the end of the dosing period, animals were exposed to different behavioural paradigms. After the behavioural tests, animals were sacrificed, the cerebral cortex was homogenised for the assessment of biochemical assays. The result showed an increase in body weight, food intake, locomotor activity, rearing and self grooming with CLOM, LETR and CLOM/LETR in all treated groups. Decreased spatial working memory and anxiolysis was observed with letrozole and/or clomiphene. Increased oxidative stress, decreased total antioxidant capacity, altered inflammatory cytokines and brain neurotransmitter were observed with letrozole and /or clomiphene. In conclusion, the administration of clomiphene and/or letrozole was associated with significant alterations in brain function, oxidative stress, inflammatory markers and brain neurotransmitter levels.

## 1.0 Introduction

Hyperandrogenism is the excessive production of male androgenic hormones, particularly in females in whom androgens are only normally present in very small quantities (Ashraf et al., 2019; Sharma and Welt, 2021; Kanasaki et al., 2022). Clinically, testosterone (which is peripherally converted to dihydrotestosterone) is considered the most important androgen in hyperandrogenism, (Sharma and Welt, 2021). In females, excessive androgen levels disrupt the hypothalamic–pituitary–gonadal axis inducing reproductive dysfunction; and presenting with clinical features that include acne, hirsuitism, obesity, amenorrhea and/or hypomenorrhea (Wang et al., 2022). In the brain, there have been reports that androgens play crucial roles in the organisation, reorganisation and programming of the circuitry (Rubinow and Schmidt, 1996; Ambar and Chiavegatto, 2009; Costine et al., 2010; Onakomaiya et al., 2014; Onaolapo et al., 2016; Onaolapo and Onaolapo, 2022). Extensive research in humans and experimental animals have shown that androgens modulate sexual behaviours, aggression, emotionality, mood, cognition and personality (Oberlander and Henderson, 2012; Onakomaiya et al., 2016; Domonkos et al., 2018; Scarth and Bjørnebekk, 2021; Kelly et al., 2022; Onaolapo and Onaolapo, 2022).

Over the last several decades there have been reports on the neurobiological, neurobehavioural, neurochemical and neuromorphological effects of excess androgen levels on the brain (Rubinow and Schmidt, 1996; Ambar and Chiavegatto, 2009; Onaolapo et al., 2016). While the psychological and behavioural impact of excessive exogenous anabolic androgenic steroid administration and increased endogenous androgen levels in males has been the subject of investigation in the last several decades (Bahrke and Yesalis, 1990; Ambar and Chiavegatto, 2009), it is only in the last decade or more that associations have begun to be made between hyperandrogenism and the development of psychiatric illness in females (Costine et al., 2009; Wang et al., 2022). Wang et al (2022) reported that using the Cox proportional analysis to compare the risk of psychiatric disorders in a cohort of women with clinical features of hyperandrogenism during a 16 year follow-up period; they observed an increased risk of psychiatric illness particularly in those aged between 20 and 29 years (Wang et al., 2022).

The management of hyperandrogenism is usually multimodal, tailored towards alleviating presenting symptoms and clinical features; with different therapies instituted for the treatment of hirsuitism, acne or infertility (ACOG Committee Opinion, 2019; Shamim et al., 2022). In women in the reproductive age group, infertility is usually a very common factor leading to a diagnosis of polycystic ovarian disease and hyperandrogenism (Messinis, 2005; Cunha and Póvoa. 2021). During the course of management, ovulation induction agents including letrozole, clomiphene citrate and gonadotropins are prescribed. Reports that compared letrozole to clomiphene citrate, revealed that letrozole was more successful at inducing ovulation and achieving live births (Banerjee Ray et al., 2012; Franik et al., 2014, 2018; Legro et al., 2014; Amer et al., 2017), however, there have also been contrary opinions (Nahid and Sirous, 2012). Also clomiphene citrate and letrozole have been used as combination therapy with reports that this results in a higher rate of ovulation compared with letrozole alone (Hajishafiha et al., 2013).

Overall, clomiphene and or letrozole therapy are generally used for the induction of ovulation and to For years, clomiphene citrate, a selective oestrogen receptor modulator was first line option of the induction of ovulation in these women, with a 70-90% success rate in inducing ovulation (Cunha and Póvoa. 2021). However, low pregnancy rates, clomiphene citrate resistance and clomiphene citrate’s anti-oestrogen effect led to research into other drugs like letrozole for the induction of ovulation s (Cunha and Póvoa. 2021). Letrozole is a non selective non-steroidal third-generation aromatase inhibitor that has been shown to be successful in inducing ovulation in women with clomiphene citrate resistance (Brown et al., 2009; Misso et al., 2012; Cunha and Póvoa. 2021). increase the chances of life births. There is ample evidence of the individual effects of letrozole or clomiphene on the brain (Aydin et al., 2008; Taylor et al., 2017; Zameer and Vohora, 2017; Gervais et al., 2019; Edwards et al., 2023), including its memory impairing and gene altering effects. While the beneficial effects of clomiphene and/or letrozole in increasing the chances of achieving pregnancy in women with hyperandrogenism are not in dispute, there is a dearth of scientific information regarding their ability to possibly alleviate (or worsen) the neurobehavioural and neurochemical effects of hyperandrogenism. Therefore this study examined the effects of clomiphene and letrozole alone or in combination in a model of hyperandrogenism in female rats.

## 2.0 Materials and Methods

### 2.1 Chemicals and drugs

Letrozole 2.5 mg tablets (Novartis Pharma), Clomiphene citrate (Clomid) 50 mg tablets, Testosterone enantate, Corn oil, Assay kits for lipid peroxidation (malondialdehyde), interleukin-10, tumour necrosis factor alpha and total antioxidant capacity (Biovision Inc., Milpitas, CA, USA).

### 2.2 Animals

Healthy weaned female Wistar rats used in this study were obtained from the animal house of the Ladoke Akintola University of Technology Ogbomoso, Oyo State, Nigeria. Rats were housed in wooden cages measuring 20 x 10 x 12 inches in temperature-controlled (22.5°C ±2.5°C) quarters with lights on at 7.00 am. Rats were allowed free access to food and water. All procedures were conducted in accordance with the approved protocols of the Ladoke Akintola University of Technology and within the provisions for animal care and use prescribed in the scientific procedures on living animals, European Council Directive (EU2010/63).

### 2.3 Diet

All animals were fed the commercially available standard rodent chow (29% protein, 11% fat, 58% carbohydrate).

### 2.4 Experimental methodology

Eighty weaned (21 day old) female rats weighing 60—70 g each were randomly assigned into eight groups of ten (n=10) animals each. Animals were grouped (Table 2.1) as normal control administered vehicle (normal saline) orally at 10 ml/kg, clomiphene (CLOM at 100 microgram/kg body weight), letrozole (LETR at 5mg/kg body weight) or combined clomiphene and letrozole groups. There were also four groups [Testosterone (TEST) control, TEST/CLOM, TEST/LETR or TEST/CLOM+LETR] of animals in which hyperandrogenism was induced with testosterone enantate administered subcutaneously at 1 mg/100 g body weight (Shirooie et al., 2021). Subcutaneous injection of testosterone or normal saline was administered daily from day 1-35, beginning on day 36, clomiphene, letrozole or saline was administered daily for 10 days. Food intake and body weight were measured using a weighing scale. At the end of the experimental period, animals were exposed to the open-field paradigm (for assessment of horizontal locomotion, rearing and self grooming behaviours), the Y maze and radial arm maze (for assessment of spatial working memory), and elevated plus maze (for the evaluation of anxiety related behaviours)and social interaction test. Twenty-four hours after the last behavioural test, animals were sacrificed by cervical dislocation, the brain was dissected, sectioned and homogenised for the assessment of inflammatory cytokines (interleukin 10, and TNF-α), lipid peroxidation (measured as malondialdehyde concentration), antioxidant status (Total antioxidant capacity) and neurotransmitter levels.

**Table 2.1.** Experimental method.

| Groups | Oral Saline | Testosterone | Subcut, saline | Neurobehaviour | Neurochemistry |
| --- | --- | --- | --- | --- | --- |
| **Control** | Yes | No | Yes | Yes | Yes |
| **CLOM** | Yes | No | Yes | Yes | Yes |
| **LETR** | Yes | No | Yes | Yes | Yes |
| **CLOM/LETR** | Yes | No | Yes | Yes | Yes |
| **TEST** | No | Yes | No | Yes | Yes |
| **TEST/CLOM** | No | Yes | No | Yes | Yes |
| **TEST/LETR** | No | Yes | No | Yes | Yes |
| **TEST/CLOM/LETR** | No | Yes | No | Yes | Yes |

Subcutaneous injection, saline: subcutaneous injection of saline

### 2.1 Assessment of body weight and food intake

Body weights of animals in all groups were measured weekly using a Mettler Toledo (Type BD6000, Switzerland) electronic weighing balance, while food consumption was assessed daily as previously described Onaolapo et al 2016).

### 2.2 Behavioural tests

Novelty induced response to the behavioural paradigms were measured by ensuring use of animals that were naïve to the paradigms as used in previous studies (Onaolapo et al., 2016, 2023) Behavioural tests were carried out in this sequence: 1) elevated plus maze, 2.) open field and 3) memory tests, 4) behavioural despair. All behavioural tests were conducted in a quiet room between the hours of 8 a.m. and 2 p.m. daily, recorded using a digital video camera and later scored by two independent observers. On each test day, rats were transported in their home cages to the behavioural testing room, and allowed to acclimatize for about 30 minutes before the behavioural tests were commenced. For all tests, approximately 5 minutes was allowed between individual rats to ensure dryness of the maze and dispersal of the residual odour of alcohol used to clean the maze as previously described Onaolapo et al. (2016). At the beginning of the behavioural tests, each animal was placed in the apparatus and its behaviour scored. After testing, each rat was removed from the maze and returned to its home cage, and all interior surfaces were cleaned thoroughly with 70 % ethanol and then wiped dry to remove any trace of odour.

#### 2.2.1 Anxiety Model: Elevated plus-maze

The elevated plus-maze (EPM) is used to measure anxiogenic or anxiolytic behaviours which are scored as time spent in the closed or open arms within a 5 minute period of exposure to the maze. Anxiety behaviours were scored as previously described (Onaolapo et al., 2014, 2017a; Olofinnade et al., 2023). The EPM is a plus-shaped apparatus with four arms that are placed at right angles to each other. The two open arms lie across from each other measuring 50 x 10 x 5 cm and perpendicular to two closed arms measuring 50 x 10 x 30 cm. The closed arms have a high wall (30 cm) to enclose the arms whereas the open arms have no side wall, only a latch that prevents animals’ from falling. Following administration of zinc or vehicle, rats were placed in the central platform facing the closed arm and their behaviour recorded for 5 minutes. The criterion for arm visit was considered only when the animal had decisively moved all its four limbs into an arm. The maze was cleaned with 70 % ethanol after each trial. The percentage of time spent in the arms was calculated as time in open arms or closed arm/total time x100, the number of entries into the arms was calculated using number of entries into open or closed arms/total number of entries.

#### 2.2.2 Open field behaviours

Exposure to the Open-field in rodents is used to measure central behaviours. Self-grooming is used to depict stereotypic behaviours. Generally central behaviours are indicative of a rat’s ability to explore open spaces. Ten minutes of open-field exposure was used to assess and score horizontal locomotion, rearing and self-grooming behaviours. The open-field paradigm is a rectangular box with a white painted hard floor that measuring 72 x 72 x 26 cm. The hard wood floor was divided by permanent red markings into 16 equal squares. The placement of animal within the box, its movement patterns and scoring are as described (Onaolapo et al., 2015, 2020a). Briefly after treatment, each rat was introduced into the field and the total locomotion (number of floor units entered with all paws), rearing frequency (number of times the animal stood on its hind legs or with its fore arms against the walls of the observation cage or free in the air) and frequency of grooming (number of body cleaning with paws, picking of the body and pubis with mouth and face washing actions) within each 10 minute interval were recorded. Behavioural tests were done between 7.00 am and 3.00 pm daily.

#### 2.2.3 Memory tests (Y-maze)

The Y-maze maze is employed for the measurement of spatial working memory. Spatial working-memory is assessed by monitoring spontaneous alternation behaviour of a rat placed in the maze for five minutes. Spontaneous alternation behaviour assesses the propensity of rodents to alternate conventionally non-reinforced choices of the Y maze on successive chances (Olofinnade et al., 2020a). The Y-maze used was composed of three equally spaced arms ((120°, 50cm long and 20 cm high, 10 cm wide). Each rat was placed in one of the arm compartments and allowed to move freely until its tail completely entered another arm. The sequence of arm entries was recorded. An alternation was defined as entry into all three arms consecutively. The number of actual alternations is number of sequential arm entries into three arms, designated A, B and C. The percentage alternation was calculated as {(Actual alternations/Total arm entry minus two) x 100} in a 5 minute interval as previously described (Onaolapo et al., 2021a; Olofinnade et al., 202ob).

### 2.3 Homogenization of the brain

Homogenates of the brain was prepared with ice-cold phosphate buffered saline using a Teflon-glass homogeniser. The homogenate was centrifuged at 5000 rev/min, 4°C, for 15 minutes. The supernatant obtained was then used for estimation of antioxidant status (Total antioxidant status and superoxide dismutase activity), lipid peroxidation levels (malondialdehyde) and inflammatory markers (Tumour necrosis factor alpha and interleukin-10).

### 2.4 Biochemical assays

#### 2.4.1 Estimation of MDA content (Lipid peroxidation)

Lipid peroxidation level was measured as MDA content. The MDA level was determined by measuring thiobarbituric acid reactive substance (Olofinnade et al., 2021a). Reactive substances of thiobarbituric acid combine with free MDA in serum to produce a coloured complex (TBAR-MDA adducts) which is then measured at an absorbance of 532 nm. The MDA concentration is expressed as μmol/L.

#### 2.4.2 Antioxidant activity

Total antioxidant capacity was measured using the trolox equivalent antioxidant capacity assay which is based on the ability of antioxidants in a sample to reduce or inhibit oxidized products generated in the assay. Total antioxidant capacity assay was assayed following previously described protocols (Olofinnade et al., 2021a; Onaolapo et al., 2022).

#### 2.4.3 Tumour necrosis factor-α and Interleukin (IL) -10

Tumour necrosis factor-α and interleukin (IL)-10 levels were measured using enzyme-linked immunosorbent assay (ELISA) techniques with commercially available kits (Biovision Inc., Milpitas, CA, USA) designed to measure the ‘total’ (bound and unbound) amount of the respective cytokines.

#### 2.4.4 Acetylcholine, dopamine, serotonin and BDNF levels

Supernatants decanted from homogenate of the hippocampus was used to assay for levels of acetylcholine (ab65345), dopamine (ab285238), serotonin (ab133053) and brain derived neurotrophic factor (using commercially available Enzyme linked immunosorbent assay kits) according to the instructions of the manufacturer. (abcam Biotechnology Company, Cambridge, United Kingdom).

### 2.7 Statistical analysis

Data were analysed with Chris Rorden’s ANOVA for windows (version 0.98). Data analysis was by One-way analysis of variance (ANOVA) and post-hoc test (Tukey’s HSD) was used for within and between group comparisons. Results were expressed as mean ± S.E.M. and p < 0.05 was taken as the accepted level of significant difference from control.

### 3.0 Result and Discussion

### 3.1 Effect of clomiphene and letrozole on body weight and food intake

Figure 1 shows the effect of clomiphene and letrozole on mean weekly body weight in a model of hyperandrogenism in female rats. There was a significant (p<0.001] increase in weekly body weight in the groups administered clomiphene (CLOM), Letrozole (LETR), CLOM/LETR, testosterone (TEST), TEST/CLOM, TEST/LET anf TEST/CLOM+LETR compared to control. Compared to TEST, weekly body weight reduced significantly with TEST/CLOM, TEST/LETR and TEST/CLOM+LETR. Analysis of relative weight change data revealed a significant (p<0.001] increase in body weight in all experimental groups compared to control. However compared to TEST, relative body weight decreased significantly with TEST/CLOM, TEST/LETR and TEST/CLOM+LETR

**Figure 1:**
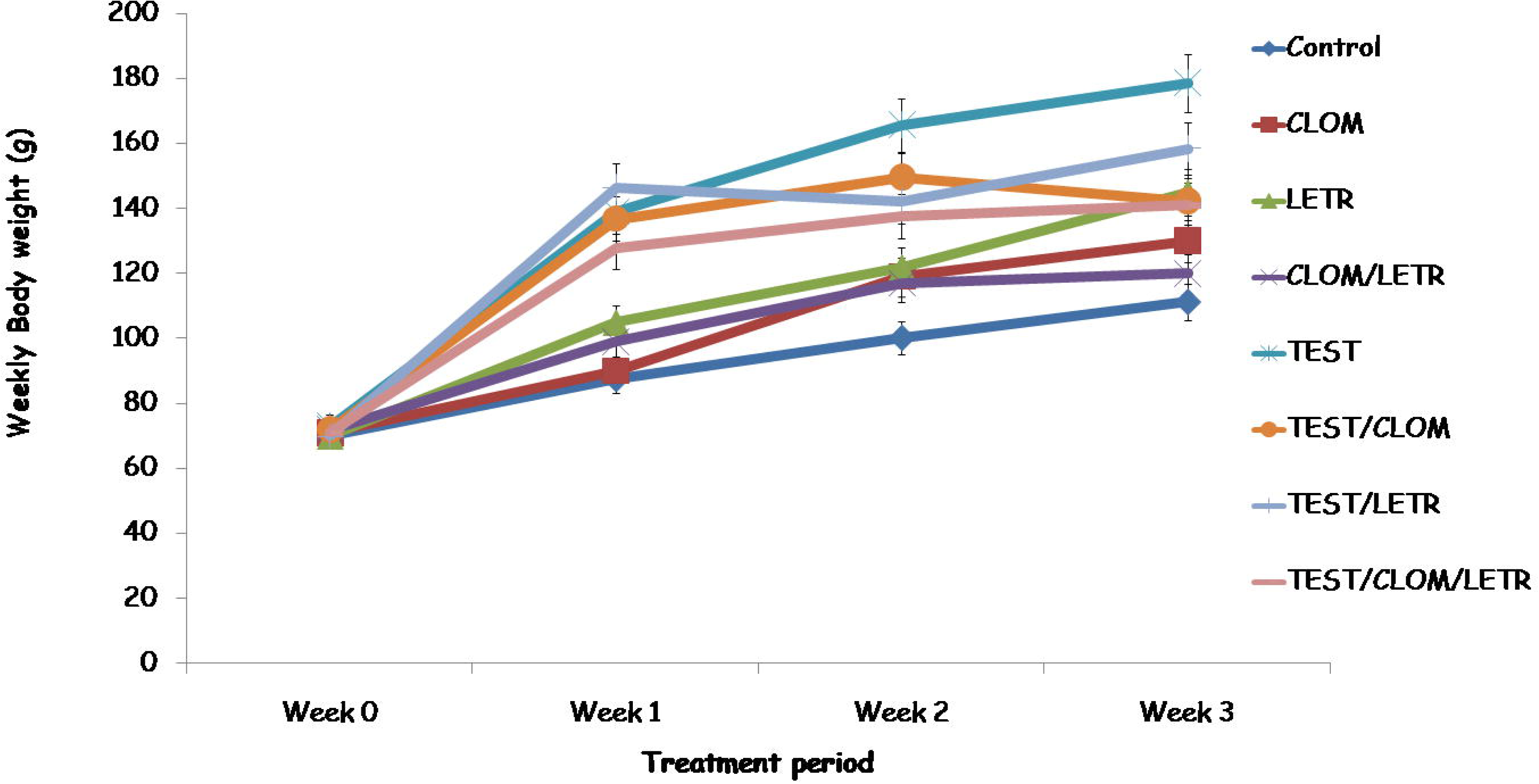
Effect of clomiphene and letrozole on weekly body weight in a model of hyperandrogenism in female rats. Each bar represents Mean ± S.E.M, number of rats per treatment group =10. TEST: Testosterone, CLOM: Clomiphene, LETR: Letrozole

Figure 2 shows the effect of clomiphene and letrozole on mean weekly food intake in a model of hyperandrogenism in female rats. There was a significant (p<0.001] increase in weekly food intake with CLOM, LETR, TEST, TEST/CLOM, TEST/LETR and a decrease with CLOM/LETR and TEST/CLOM/LETR compared to control. Compared to TEST, weekly food intake reduced significantly with TEST/CLOM, TEST/LETR and TEST/CLOM+LETR.

**Figure 2:**
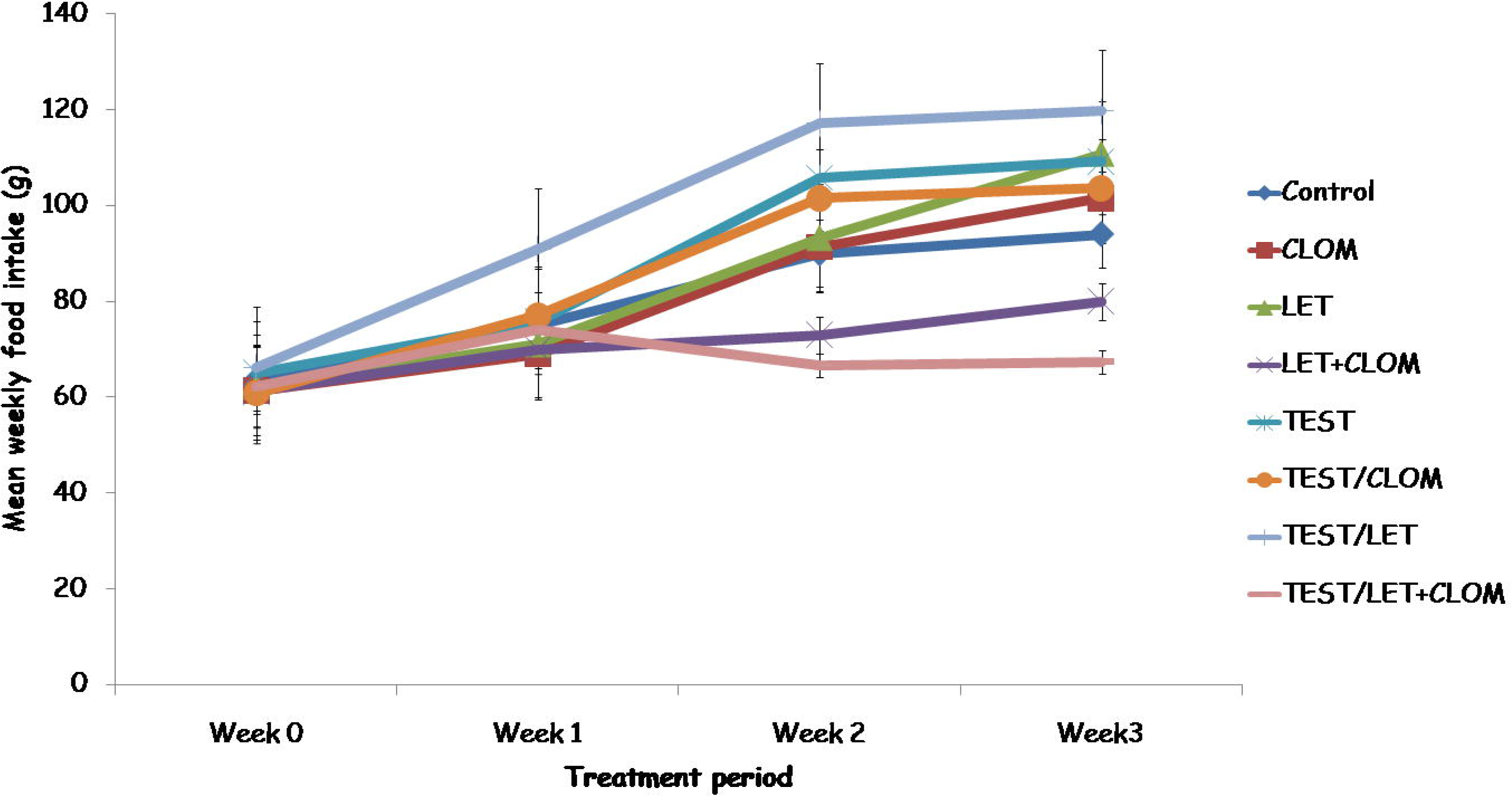
Effect of clomiphene and letrozole on weekly food intake in a model of hyperandrogenism in female rats. Each bar represents Mean ± S.E.M, number of rats per treatment group =10. TEST: Testosterone, CLOM: Clomiphene, LETR: Letrozole.

### 4.2 Effect of clomiphene and letrozole on open field exploratory behaviours

Figure 3 shows the effect of clomiphene and letrozole on open field exploratory behaviours including horizontal locomotion (upper panel) and vertical locomotion (lower panel) in a model of hyperandrogenism in female rats. Horizontal locomotion measured as line crossing decreased significantly (p<0.001] with CLOM, LETR and CLOM/LETR and increased significantly with TEST, TEST/CLOM, TEST/LETR and TEST/CLOM+LETR compared to control. Compared to TEST, horizontal locomotion increased significantly with TEST/CLOM, TEST/LETR and TEST/CLOM+LETR. Vertical locomotion measured as rearing decreased significantly (p<0.001] with CLOM, LETR and CLOM/LETR and increased significantly with TEST, TEST/CLOM, TEST/LETR and TEST/CLOM+LETR compared to control. Compared to TEST, rearing activity increased significantly with TEST/CLOM, TEST/LETR and TEST/CLOM+LETR.

**Figure 3:**
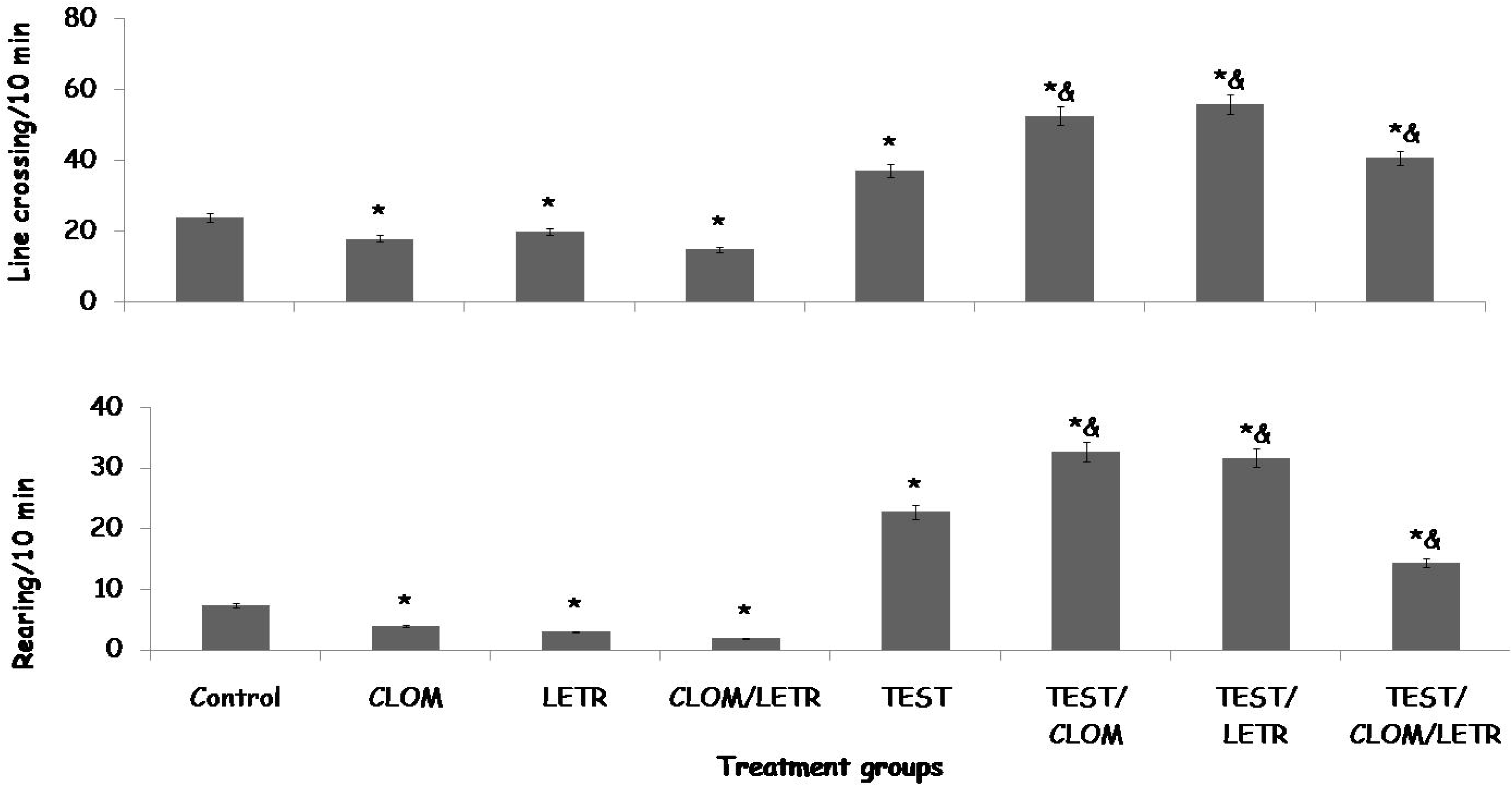
Effect of clomiphene and letrozole on line crossing (upper panel) and rearing (lower panel) in a model of hyperandrogenism in female rats. Each bar represents Mean ± S.E.M, number of rats per treatment group =10. TEST: Testosterone, CLOM: Clomiphene, LETR: Letrozole.

### 4.3 Effect of clomiphene and letrozole on open field self-grooming behaviour and spatial working memory in the Y maze

Figure 4 shows the effect of clomiphene and letrozole on open field self grooming behaviour (upper panel) and Y-maze spatial working memory (lower panel) in a model of hyperandrogenism in female rats. Self grooming decreased significantly (p<0.001] with CLOM, LETR, CLOM/LETR and TEST, while it increased significantly with TEST/CLOM, TEST/LETR and TEST/CLOM+LETR compared to control. Compared to TEST, self grooming increased significantly with TEST/CLOM, TEST/LETR and TEST/CLOM+LETR.

**Figure 4:**
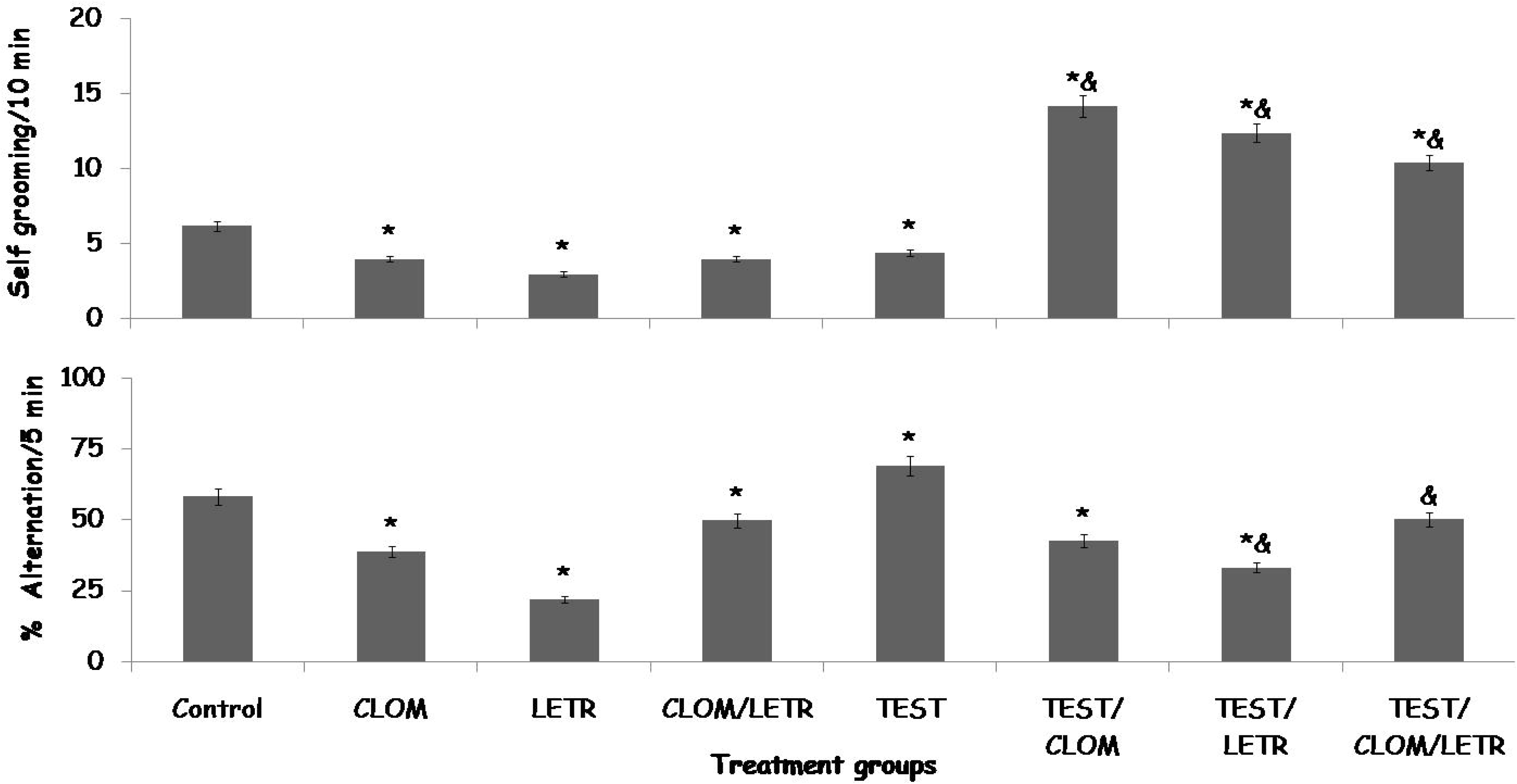
Effect of clomiphene and letrozole on self grooming (upper panel) and Y-maze spatial working memory (lower panel) in a model of hyperandrogenism in female rats. Each bar represents Mean ± S.E.M, number of rats per treatment group =10. TEST: Testosterone, CLOM: Clomiphene, LETR: Letrozole.

Y maze spatial working memory measured as percentage alternation decreased significantly (p<0.001] with CLOM, LETR. CLOM/LETR, TEST/CLO and TEST/LETR while it increased significantly with TEST compared to control. Compared to TEST control, Y maze spatial working memory decreased significantly in groups treated with TEST/CLOM, TEST/LETR and TEST/CLOM+LETR.

### 4.4 Effect of clomiphene and letrozole on anxiety related behaviours

Figure 5 shows the effect of clomiphene and letrozole on open arm time (upper panel) and closed arm time (lower panel) in a model of hyperandrogenism in female rats. Anxiety behaviour measured using open arm time and closed arm time. Open arm time decreased significantly (p<0.001] with CLOM, LETR and CLOM/LETR, while an increase was observed with TEST, TEST/CLOM, TEST/LETR and TEST/CLOM+LETR compared to control. Compared to TEST, open arm time increased significantly with TEST/CLOM, TEST/LETR and TEST/CLOM+LETR.

**Figure 5:**
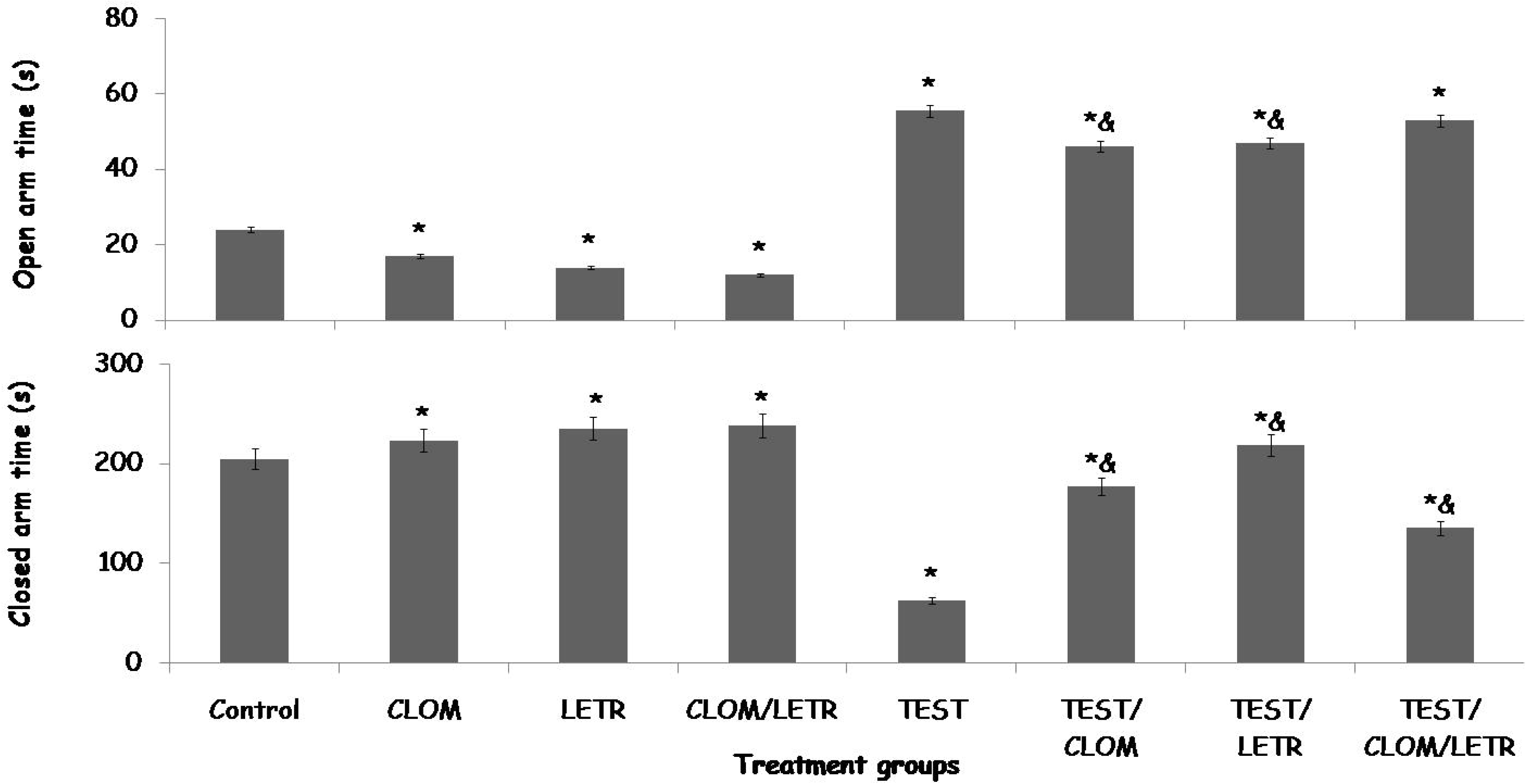
Effect of clomiphene and letrozole on open arm time (upper panel) and closed arm time (lower panel) in a model of hyperandrogenism in female rats. Each bar represents Mean ± S.E.M, number of rats per treatment group=10. TEST: Testosterone, CLOM: Clomiphene, LETR: Letrozole.

Closed arm time increased significantly (p<0.001] with CLOM, LETR and CLOM/LETR, and decreased significantly with TEST, TEST/CLOM, TEST/LETR and TEST/CLOM+LETR compared to control. Compared to TEST, closed-arm time increased significantly in groups treated with TEST/CLOM, TEST/LETR and TEST/CLOM+LETR.

### 4.5 Effect of clomiphene and letrozole on lipid peroxidation, Total antioxidant capacity and gonadotropin levels

Table 1 shows the effect of clomiphene and letrozole on lipid peroxidation, Total antioxidant capacity and gonadotropin levels in testosterone treated female rats. Lipid peroxidation measured as malondialdehyde (MDA) levels increased significantly with CLOM, LETR, CLOM/LETR, TEST, TEST/CLOM, TEST/LETR and TEST/CLOM+LETR compared to control. Compared to TEST, MDA levels increased significantly with TEST/CLOM, TEST/LETR and decreased with TEST/CLOM+LETR. Total antioxidant capacity (TAC) decreased significantly with CLOM, LETR, CLOM/LETR, TEST, TEST/CLOM, TEST/LETR and TEST/CLOM+LETR compared to control. Compared to TEST, TAC decreased significantly with TEST/CLOM, TEST/LETR and increased with TEST/CLOM+LETR.

**Table 1:**
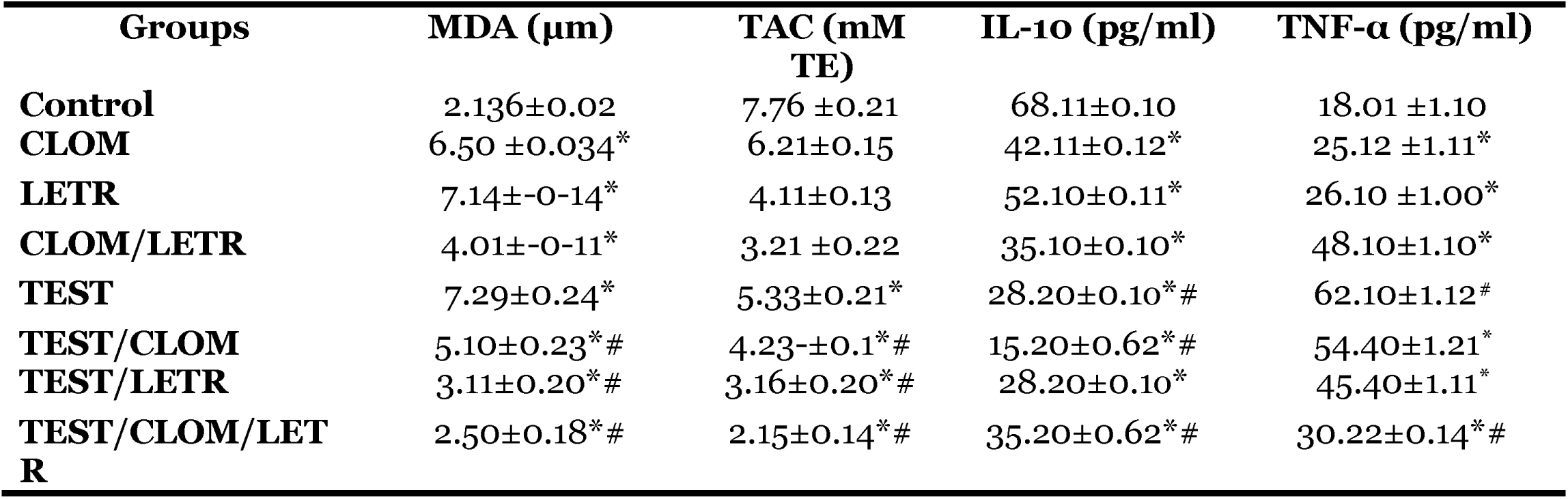
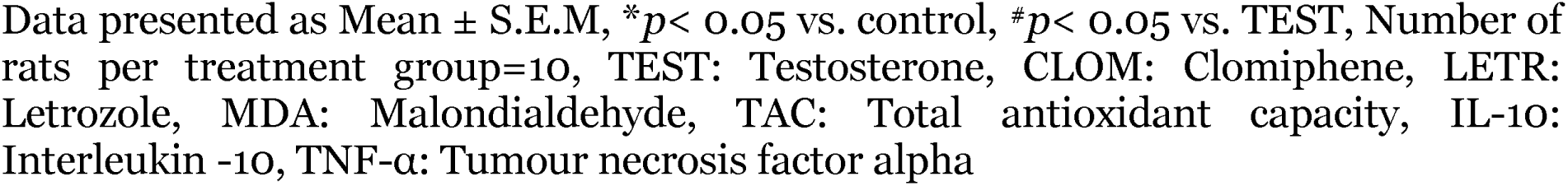
Effect of clomiphene and letrozole on lipid peroxidation, Total antioxidant capacity and inflammatory markers.

Interleukin-10 (IL-10) levels decreased significantly with CLOM, LETR, CLOM/LETR, TEST, TEST/CLOM, TEST/LETR and TEST/CLOM+LETR compared to control. Compared to TEST, IL-10 levels decreased significantly in groups administered TEST/CLOM, and increased with TEST/CLOM+LETR.

Tumour necrosis factor (TNF-α) levels increased significantly with CLOM, LETR, CLOM/LETR, TEST, TEST/CLOM, TEST/LETR and TEST/CLOM+LETR compared to control. Compared to TEST, TNF-α levels decreased significantly with TEST/CLOM, TEST/LETR and TEST/CLOM+LETR.

### 4.7 Effect of clomiphene and letrozole on brain neurotransmitter levels

Table 2 shows the effect of clomiphene and letrozole on neurotransmitter levels in the brain of testosterone treated female rats. Dopamine and acetylcholine levels decreased significantly with CLOM, LETR, CLOM/LETR, TEST, TEST/CLOM and TEST/LETR compared to control. Compared to TEST, dopamine and acetylcholine levels increased significantly with TEST/CLOM, TEST/LETR and TEST/CLOM+LETR.

**Table 2:**
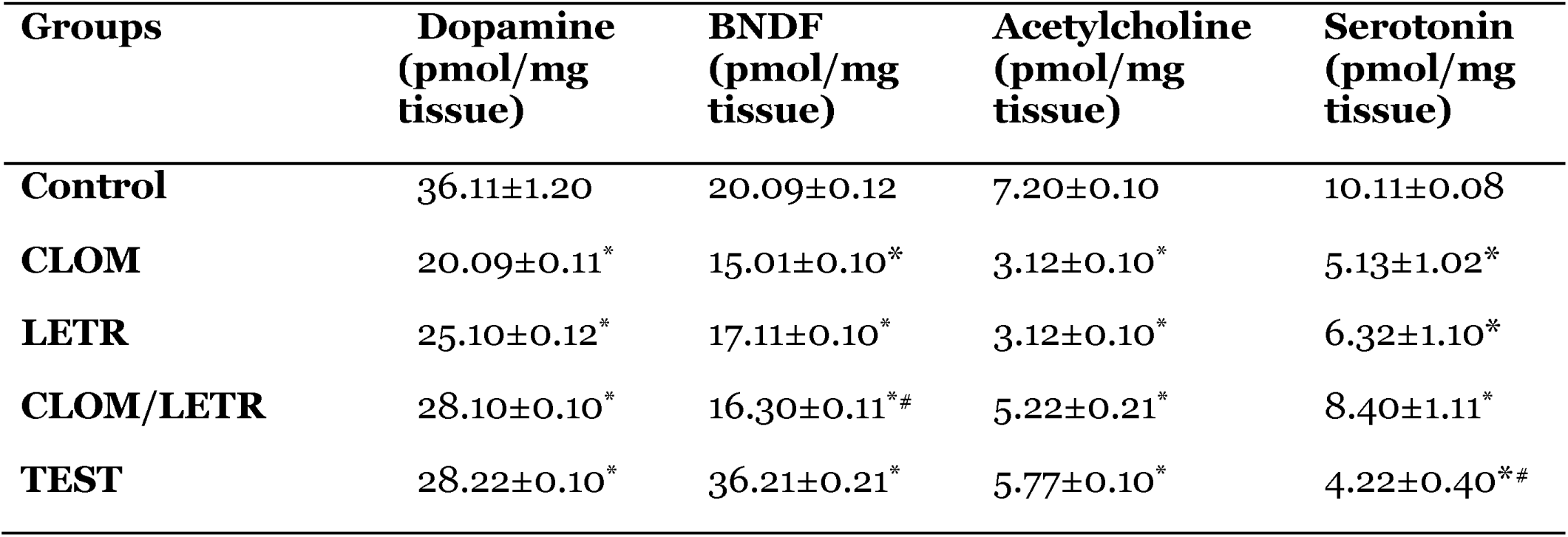
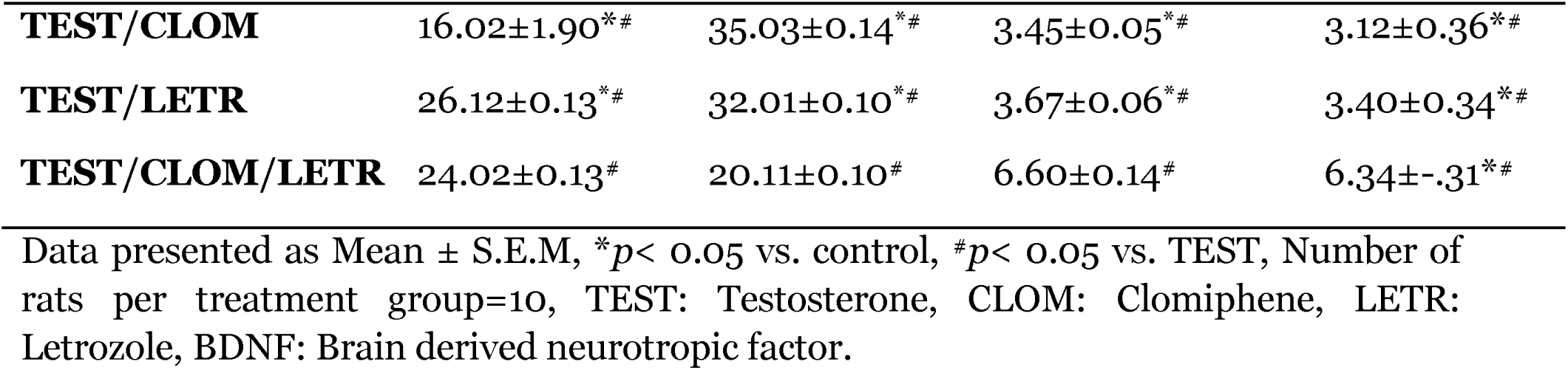
Effect of clomiphene and letrozole on neurotransmitter levels.

| Groups | Dopamine<br>(pmol/mg<br>tissue) | BNDF<br>(pmol/mg<br>tissue) | Acetylcholine<br>(pmol/mg<br>tissue) | Serotonin<br>(pmol/mg<br>tissue) |
| --- | --- | --- | --- | --- |
| **Control** | 36.11±1.20 | 20.09±0.12 | 7.20±0.10 | 10.11±0.08 |
| **CLOM** | 20.09±0.11* | 15.01±0.10* | 3.12±0.10* | 5.13±1.02* |
| **LETR** | 25.10±0.12* | 17.11±0.10* | 3.12±0.10* | 6.32±1.10* |
| **CLOM/LETR** | 28.10±0.10* | 16.30±0.11*# | 5.22±0.21* | 8.40±1.11* |
| **TEST** | 28.22±0.10* | 36.21±0.21* | 5.77±0.10* | 4.22±0.40*# |
| **TEST/CLOM** | 16.02±1.90*<sup>#</sup> | 35.03±0.14*<sup>#</sup> | 3.45±0.05*<sup>#</sup> | 3.12±0.36*<sup>#</sup> |
| **TEST/LETR** | 26.12±0.13*<sup>#</sup> | 32.01±0.10*<sup>#</sup> | 3.67±0.06*<sup>#</sup> | 3.40±0.34*<sup>#</sup> |
| **TEST/CLOM/LETR** | 24.02±0.13<sup>#</sup> | 20.11±0.10<sup>#</sup> | 6.60±0.14<sup>#</sup> | 6.34±.31*<sup>#</sup> |

Data presented as Mean ± S.E.M, \* $p < 0.05$ vs. control, <sup>#</sup> $p < 0.05$ vs. TEST, Number of rats per treatment group=10, TEST: Testosterone, CLOM: Clomiphene, LETR: Letrozole, BDNF: Brain derived neurotropic factor.

Brain derived neurotropic factor (BDNF) levels decreased significantly with CLOM, LETR, CLOM/LETR and increased with TEST, TEST/CLOM and TEST/LETR compared to control. Compared to TEST, BDNF decreased significantly with TEST/CLOM, TEST/LETR and TEST/CLOM+LETR.

Serotonin levels decreased significantly with CLOM, LETR, CLOM/LETR, TEST, TEST/CLOM, TEST/LETR and TEST/CLOM+LETR compared to control. Compared to TEST, serotonin levels increased significantly with TEST/CLOM, TEST/LETR and TEST/CLOM+LETR.

## Discussion

This study examined the effects of clomiphene citrate and/or letrozole on body weight, neurobehaviour, antioxidant status, inflammatory response and brain neurotransmitter levels in testosterone induced hyperandrogenism in female rats. The result showed an increase in body weight, food intake, locomotor activity, rearing and self grooming in the healthy animals and the treated groups with testosterone induced-hyperandrogenism. Spatial working memory and anxiolysis increased with hyperandrogenism and decreased with letrozole and/or clomiphene. Increased oxidative stress, decreased total antioxidant capacity, altered inflammatory cytokines and brain neurotransmitter were observed with letrozole and/or clomiphene administered in healthy rats and in animals with testosterone induced hyperandrogenism. However which was mildly reversed with clomiphene and/or letrozole treatment.

It is widely accepted that hyperandrogenism in females is closely related to obesity, with reports that weight gain could also contribute significantly towards worsening of hyperandrogenism, hampering efforts to establish effective weight control (Barber et al., 2019). In this study, testosterone-induced hyperandrogenism was associated with weight gain and increased food intake in female rats. Results of a previous study from our laboratory that had evaluated age-related effects of testosterone administration in gonadally-intact males revealed that the subcutaneous injection of testosterone was associated with weight gain (Onaolapo et al., 2016). In this study also the administration of testosterone enantate to weaned female rats was associated with weight gain and metabolic derangement (data not included) which are consistent with PCOS as previously described by Shirooie et al. (2021). In the healthy female rats administered clomiphene and/or letrozole a gradual increase in body weight was observed while the co-administration of clomiphene and letrozole was associated with lesser weight gain compared to clomiphene or letrozole alone. In the groups with testosterone induced hyperandrogenism, treatment resulted in an increase in weight in groups treated with letrozole and a decrease in weight in the groups treated with clomiphene either alone or when co-administered with letrozole, when compared to healthy control; although overall a mitigation of testosterone-induced weight gain was observed in the three treatment groups.

Food intake increased with clomiphene and letrozole when administered alone compared to control or testosterone control. However in the hyperandrogenism groups treated with both clomiphene and letrozole a decrease in food intake was observed. Although there is a dearth of scientific information on the possible effects of clomiphene and or letrozole on body weight and or food intake, there have been suggestions that weight gain and increased appetite are side effects of clomiphene or letrozole therapy in humans.

For decades there have been reports on the brain effects of clomiphene (Olcese et al. 1984; McMillan-Castanares et al., 2023), and letrozole (Chang et al., 2013; Gervais et al., 2019). There have been suggestions that clomiphene and/or letrozole could adversely modulate neurobehaviour and brain neurotransmitters levels. In this study administration of clomiphene or letrozole alone or in combination to healthy female rats reduced locomotor activity, rearing and grooming behaviours demonstrating a general central inhibitory effect on the brain of these rats. The quantification of physiological behaviours including locomotor activity, rearing and self-grooming behaviours have become valuable approaches for the assessment and the understanding the biological systems and neural mechanisms that underlie normal and abnormal behaviours in humans and rodents (Onaolapo and Onaolapo et al., 2013; Minassian et al., 2016; Young et al., 2016; Onaolapo et al., 2017b, 2017c). Locomotion is the spontaneous movement of an animal in all direction. It is modulated through the activities of neurotransmitters (dopamine, glutamate and even oestrogen) in the brain. Oestrogen has been shown to increase locomotor activity, Ogawa et al. (2003) reported that oestrogen had the ability to increase running wheel activity in rodents through its actions on the medial preoptic area of the brain. In another study, Espinosa and Curtis, (2018) reported that oestrogen related increase in locomotion was related to an increase in the levels of dopamine in the nucleus accumbens. The effects observed in this study following administration of anti-oestrogen clomiphene which resulted in a decrease locomotion rearing and grooming would suggests that oestrogens affect open field behaviour generally. Also, observed was a decrease in cerebral cortex levels of dopamine which also support the results of the study by Espinosa and Curtis, (2018). Aromatase is an enzyme that converts androgens to oestrogen or oestradiol and is also important in the militating the effects of sex hormones on the brain (Borbélyová et al., 2017). In groups administered aromatase inhibitor letrozole, a decrease in open field behaviours was also observed. This is would suggest that while letrozole does have anti-oestrogen effects, the effect of oestrogen on open field behaviour observed in this study could be attributed to the effects of oestrogen converted from androgens through the action of aromatase on androgens and not necessarily the oestrogens secreted de novo in the brain. In the testosterone treated rats (hyperandrogenism model) an increase in horizontal locomotion and rearing and a decrease in self-grooming was observed. Testosterone has been shown modulate behavioural function (Onaolapo et al., 2016,) effects that are mediated through its effects on different receptors. In a study from our laboratory examining the impact of age on testosterone effect in male rodents, a decrease in locomotor activity, rearing and grooming was observed in prepubertal male rats (similar age to female rats used in this study) after 21 days of administration of testosterone. This would suggest that while in males testosterone was associated with a central inhibitory effect, in female rats’ testosterone administration was associated with a central excitatory effect. The central excitatory effects observed with horizontal locomotion and rearing could be linked to the possible aromatization of testosterone to oestrogen in these female rats. In the hyperandrogenism groups treated with clomiphene or letrozole an increase in locomotion, rearing and grooming was observed compared to the testosterone control group, however in the group treated with a combination of clomiphene and letrozole a decrease in open-field behaviours was observed compared to either the clomiphene or letrozole alone groups. While there is a dearth of scientific literature on the effects of clomiphene and letrozole on the brain the results of this study suggests that the effect observed with the combination is possibly a resultant of the neurotransmitter effects of clomiphene, letrozole and testosterone. The behavioural changes observed following combination of clomiphene and letrozole could also be attributed to the effects of luteinizing hormone or follicle stimulating hormone, Studies have shown that both clomiphene and letrozole on their own result in the release of gonadotropins (Which was also observed in this study). Letrozole causes a decrease in oestrogen levels in the brain thereby stimulating the brain to increase gonadotropins secretion; clomiphene on the other hand lowers the negative feedback of oestrogen, which eventually leads to an increase in pituitary gonadotropin hormones (Hajishafiha et al., 2013). Luteinizing hormone has also been shown to have effects on brain function (Mora and Díaz-Véliz, 1983; Saito, 1988; de Beun et al., 1991).

The effect of steroid hormones on cognition and anxiety related behaviours have been reported (Celec et al., 2015; Hara et al.,2015; Domonkos et al., 2018; Renczés et al., 2020) . There have been reports that oestrogen facilitates cognitive function through its effects on oestrogen receptors located in the prefrontal cortex and hippocampus (Hara et al., 2015). Oestradiol has also been shown to have an anxiolytic effect (Renczés et al., 2020). In this study the administration of clomiphene and/or letrozole was associated with a decrease in spatial working memory in all groups except the testosterone control group. The results of this study support the result of studies that have demonstrated the memory impairing effects of clomiphene and/or letrozole (Gervais et al., 2019; Marbouti et al., 2020). Although there is a dearth of scientific information on the memory impairing effects of clomiphene either alone or in combination with letrozole, the memory impairing effects of letrozole have been attributed to its ability to increase hippocampal dopamine levels (although a decrease in cerebral cortex levels of dopamine was observed in healthy animal in this study), decrease estradiol synthesis, mitochondrial volume, synaptophysin reactivity and dendritic spine density (Chang et al., 2013; Gervais et al., 2019; Marbouti et al., 2020; Moustafa et al., 2022). Also in this study administration of clomiphene or letrozole alone or in combination to healthy rats was associated with an anxiogenic effect. While an anxiogenic response has been reported following the administration of letrozole in middle aged male rats, the results of our study contradicts the effect observed in female rats (Borbélyová et al., 2017), although the differences observed in the results could be attributable to the age of the animals, since age and sex have been reported to drug effects on the brain (Borbélyová et al., 2017). The anxiogenic effects of clomiphene observed in this study could be attributed to its anti-oestrogen effects considering that studies had reported the anxiolytic effects of oestrogen (Renczés et al., 2020). The combination of clomiphene and letrozole was associated with an increase in the levels of luteinizing hormone (LH) in this study which could be responsible for the memory impairing and anxiogenic effects observed, there have been reports of the memory impairing and anxiogenic effects of LH (Arfa-Fatollahkhani et al., 2017). In the rats with testosterone induced hyperandrogenism, improved spatial working memory and anxiolysis was observed, corroborating the results of a number of studies that have reported the anxiolytic and memory enhancing effects of testosterone (Hodosy et al., 2012; Domonkos et al., 2018). In the hyperandrogenism groups treated with clomiphene, letrozole or a combination of both drugs, a decrease in memory was observed with clomiphene and letrozole alone while in the group administered clomiphene and letrozole no significant difference in memory scores was observed compared to control and a decrease compared to the hyperandrogenism control. While there is limited scientific information on the possible brain effects of combining clomiphene and letrozole in this study we observed a decrease in cerebral cortex levels of dopamine, acetylcholine and brain derived neurotrophic factor which are neurotransmitters that have been shown to modulate brain function. The role of oxidative stress, antioxidant status and inflammatory markers in modulating brain function has also been reported (Salim, 2017; Singh et al., 2019; Onaolapo et al., 2021, 2022b). In this study the administration of clomiphene or letrozole alone or combined to healthy female rats or female rats with hyperandrogenism was associated with increased lipid peroxidation and tumour necrosis factor-alpha while a decrease in total antioxidant capacity and interleukin-10. A number of studies have reported that clomiphene or letrozole were associated with increased oxidative stress and decreased antioxidant status parameters peripherally, being linked to the development of polycystic ovarian syndrome in females (Peker et al., 2021). Increased Oxidative stress and reduced levels of antiinflammatory cytokines in rats administered clomiphene and/or letrozole (healthy rats) would suggest that this could be responsible for memory impairment, anxiety and central inhibitory response in the open field. The combination of clomiphene and letrozole showed worsening of the oxidative stress and pro-inflammatory response in these groups of animals.

Just as sex hormones have been shown to significantly impact brain development and growth (Barth et al., 2015; Brann et al., 2022), ovulation induction agents like letrozole or clomiphene have been shown to alter brain neurotransmitters levels and activity (Aydin et al., 2008; Marbouti et al., 2020). Neurotransmitters and modulators like dopamine, serotonin and BDNF have been reported to modulate neuronal development, neurotransmission, and brain plasticity, strongly influencing brain behaviour, aging and cognition (Peters et al., 2021; Bazzari and Bazzari, 2022). In this study the administration of clomiphene, letrozole alone or combined was associated with a decrease in the levels of dopamine, serotonin and BDNF when compared to rats in the control group. The effects 0bserved with dopamine in this study is consistent with the results of studies examining the mechanisms by which aromatase inhibitors like letrozole cause cognitive deficits that had reported a decrease in the brain levels of catecholamines including dopamine (Aydin et al., 2008). Although there is a dearth of scientific information on the effect of clomiphene and/or letrozole on brain in healthy animals or in a background of hyperandrogenism, the reduction of neuromodulator and neurotransmitter levels could be responsible fior the neurobehavioural changes observed in this study (Bathina et al., 2015; Mariga et al., 2017; Miranda et al., 2019)

## Conclusion

In conclusion, the administration of clomiphene and/or letrozole was associated with significant alterations in brain function, oxidative stress, inflammatory markers and brain neurotransmitter levels.

## Notes

### Competing Interest Statement

The authors have declared no competing interest.

